# Delayed global feedback in the genesis and stability of spatiotemporal patterns in paced biological excitable media

**DOI:** 10.1101/2020.05.13.094011

**Authors:** Zhen Song, Zhilin Qu

**Author notes:** Correspondence to: Zhen Song, PhD, Department of Medicine, Division of Cardiology, David Geffen School of Medicine at UCLA, A2-237 CHS, 650 Charles E. Young Drive South, Los Angeles, CA 90095, Zhilin Qu, PhD, Department of Medicine, Division of Cardiology, David Geffen School of Medicine at UCLA, A2-237 CHS, 650 Charles E. Young Drive South, Los Angeles, CA 90095.

## Abstract

A multi-scale approach was used to investigate the roles of delayed global feedback (DGF) in the genesis and stability of spatiotemporal patterns in periodically-paced excitable media. Patterns that are temporal period-2 (P2) and spatially concordant (in-phase) or discordant (out-of-phase) were investigated. First, simulations were carried out using a generic spatiotemporal model composed of coupled FitzHugh-Nagumo units with DGF. When DGF is absent, concordant and discordant P2 patterns occur depending on initial conditions. The discordant P2 patterns are spatially random. When the DGF is negative, only concordant P2 patterns exist. When the DGF is positive, both concordant and discordant P2 patterns can occur. The discordant P2 patterns are still spatially random, but they satisfy that the global signal exhibits a temporal period-1 behavior. Second, to validate the spatiotemporal dynamics in a biological system, simulations were carried out using a 3-dimensional physiologically detailed ventricular myocyte model. This model can well capture the intracellular calcium release patterns widely observed in experiments. The properties of DGF were altered by changing ionic currents or clamping voltage. The spatiotemporal pattern dynamics of calcium release in this model match precisely with those of the generic model. Finally, theoretical analyses were carried out using a coupled map lattice model with DGF, which reveals the instabilities and bifurcations leading to the spatiotemporal dynamics and provides a general mechanistic understanding of the role of DGF in the genesis, selection, and stability of spatiotemporal patterns in paced excitable media.

**Author Summary:** Understanding the mechanisms of pattern formation in biological systems is of great importance. Here we investigate the dynamical mechanisms by which delayed global feedback affects pattern formation and stability in periodically-paced biological excitable media, such as cardiac or neural cells and tissue. We focus on the formation and stability of the temporal period-2 and spatially in-phase and out-of-phase patterns. Using a multi-scale modeling approach, we show that when the delayed global feedback is negative, only the spatially in-phase patterns are stable; when the feedback is positive, both spatially in-phase and out-of-phase patterns are stable. Also, under the positive feedback, the out-of-phase patterns are spatially random but satisfy that the global signals are temporal period-1 solutions.

## Introduction

Pattern formation is ubiquitous in biological systems, ranging from biological development [1, 2], ecosystems [3], to disease development [4]. Many of the pattern formation processes can be explained by Turing instability in reaction-diffusion (or activator-inhibitor) systems [5, 6]. However, pattern formation via other mechanisms has also been proposed, in particular for spatiotemporal patterns, which are also widely observed in biological systems [7–12]. The fundamental processes causing temporal and spatiotemporal dynamics in biological systems are positive and negative feedback loops [5, 6, 13]. While many studies investigated the roles of local and instantaneous feedback loops in pattern formation, studies have also carried out to investigate the roles of instantaneous global feedback and time delay global feedback (DGF) loops, such as the ones in oscillatory media of chemical reactions [14–19]. In this study, we focus on the roles of DGF in pattern formation in a class of biological systems, i.e., excitable media subjected to periodic global stimulation.

Many biological systems are excitable media with DGF loops that are not as explicit as those implemented in the chemical reaction experiments [17, 19]. Here we use intracellular calcium (Ca^2+^) signaling, which is required for many biological functions [20, 21], as an example to explain the existence of DGF. The fundamental unit of Ca^2+^ signaling in cells is called Ca^2+^ release unit (CRU) (Fig.1a). Ca^2+^ entering the cell from the voltage-gated Ca^2+^ channels triggers the opening of the Ca^2+^ release channels to release Ca^2+^ from the internal Ca^2+^ stores. The open probability of the Ca^2+^ release channels is further enhanced by the released Ca^2+^. This process is known as Ca^2+^-induced Ca^2+^ release (CICR), which is an instantaneous local feedback loop responsible for a rich spectrum of Ca^2+^ dynamics widely observed in biological systems [22–26]. Besides this instantaneous feedback loop, implicit delayed feedback loops exist, i.e., Ca^2+^ in the present beat may affect itself in the next beat (Fig.1b). This feedback can be mediated by the Ca^2+^ current (I_Ca_) of the voltage-gated Ca^2+^ channels or the Ca^2+^ release properties of the internal stores through either voltage or Ca^2+^-dependent signaling pathways. For example, in cardiac myocytes, Ca^2+^ is coupled to voltage via Ca^+^-dependent ion channels and pumps. Changing Ca^2+^ in the present beat changes the action potential duration (APD) and thus the diastolic interval (DI), affecting the recovery of voltage-gated Ca^2+^ channels in the next beat. As a result, the change in the recovery alters I_Ca_ and hence Ca^2+^, forming a delayed feedback loop. Note that in excitable cells, ion channels generally remain in closed or inactivation states in the quiescent phase. Therefore, the effects of this delayed feedback are manifested in the next beat. In other words, the time delay of the feedback loop is simply the pacing period T.

**Fig.1.**
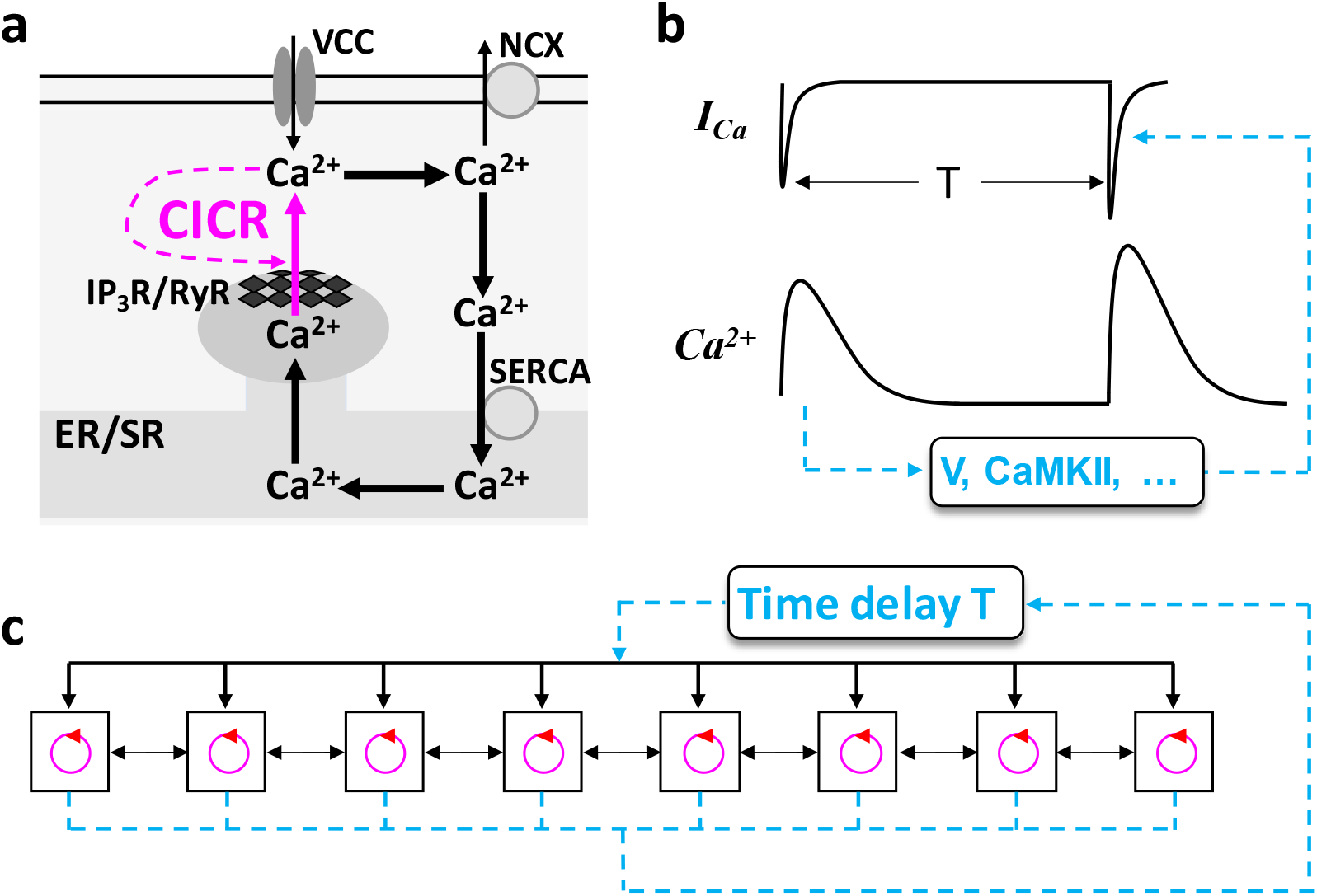
Schematic diagrams of Ca^2+^ cycling and a generic model of coupled excitable units with DGF. **a**. Schematic diagram of a basic Ca^2+^ release unit (CRU) in excitable cells. Ca^2+^ from voltage gated Ca^2+^ channels (VCCs) triggers the opening of the inositol trisphosphate receptors (IP3Rs) or ryanodine receptors (RyRs), releasing the Ca^2+^ stored in the endoplasmic or sarcoplasmic reticulum (ER/SR). The released Ca^2+^ further triggers more IP3Rs/RyRs to open, forming a positive feedback loop. This process is called Ca^2+^-induced Ca^2+^ release (CICR). Ca^2+^ is extruded by Na^+^-Ca^2+^ exchange (NCX) or other Ca^2+^ pumps and uptaken back into the ER/SR via sarco/endoplasmic reticulum Ca^2+^ ATPase (SERCA). **b**. Schematic diagram of delayed feedback in Ca^2+^ signaling via Ca^2+^ current (I_Ca_). T is the pacing period. **c**. Schematic diagram of a generic model of coupled excitable units (e.g., CRUs) with a DGF loop of time delay T.

A cell consists of thousands of CRUs which are coupled via Ca^2+^ diffusion. The CRUs are themselves excitable units [24, 27–29], which are triggered by a global signal, i.e., voltage. Therefore, one can simplify the Ca^2+^ signaling system into a coupled array of excitable units under a global stimulation with a DGF loop (Fig.1c). Since voltage is the global signal, under normal conditions, depolarization of the cell synchronizes the firings of the CRUs, resulting in a synchronous whole-cell Ca^2+^ release, such as Ca^2+^ release in neurons (Fig.S1a) [30]. The synchronous Ca^2+^ release is essential for muscle contraction [21] and many other types of biological functions [20]. However, under abnormal or diseased conditions, dyssynchronous Ca^2+^ releases can occur, such as spatially discordant Ca^2+^ alternans widely observed in cardiac myocytes (Fig.S1b) [31–33]. Although it is clear that voltage serves as the global signal to synchronize the CRU releases, it is unclear how dyssynchronous patterns are formed and what are the roles of the DGF in maintaining the synchronous release patterns or the development of the dyssynchronous release patterns.

In addition to intracellular Ca^2+^ signaling, other biological systems can also be described by the simplified scheme in Fig.1c, such as the excitation dynamics in cardiac muscle or neural networks. In cardiac tissue, myocytes are electrically excitable units that are coupled via gap junctions. Contraction of the heart can serve as the global signal, which may mediate DGF via mechano-electric feedback through activating mechano-sensitive channels and affecting intracellular Ca^2+^ release [34–36]. This DGF may play essential roles in arrhythmogenic pattern formation in the heart, such as the widely observed spatially discordant APD alternans [37, 38]. In neural networks, the roles of delayed feedback in neural firing dynamics have been investigated [39, 40], and DGF may also play essential roles in the formation and stability of clustered firing of neurons [41].

This study was set to investigate the roles of DGF in the genesis and stability of spatiotemporal patterns in periodically-paced biological excitable media, focusing on temporal period-2 (P2) and spatially concordant (in-phase) or discordant (out-of-phase) patterns. A multi-scale approach was applied. First, a generic model consisting of a coupled array of excitable units described by the FitzHugh-Nagumo (FHN) model was used, and simulations were carried out to reveal the pattern dynamics caused by DGF. To validate the findings of the generic model, we used a 3-dimensional (3D) ventricular myocyte model and carried out simulations to investigate the roles of DGF in spatially concordant and discordant Ca^2+^ alternans dynamics. Of note, the term “alternans” in the context of the cardiac systems refers to a P2 state. Finally, a coupled map lattice (CML) model was used to perform detailed theoretical analyses, which provide a general mechanistic understanding of the roles of DGF in pattern formation, selection, and stability in periodically-paced biological excitable media.

## Results

### DGF in the genesis of spatiotemporal dynamics in an array of coupled FHN units

We used a generic model consisting of a one-dimensional (1D) array of coupled FHN units to investigate the spatiotemporal excitation patterns. The governing differential equations are:

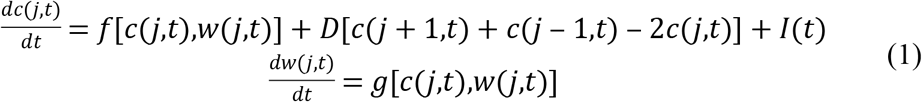

in which *j* ∈ *{1, 2, …, L},* is the spatial index of the FHN units with *L* being the length of the 1D array. We used the standard FHN kinetics, i.e.,

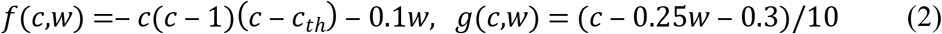

where *c* is the activator and *w* is the inhibitor. *c_th_*=0.5 is a parameter determining the threshold for excitation, and *D*=0.1 is the coupling strength. No-flux boundary condition was used. *I*(t) is the external stimulus pulse, which is formulated as

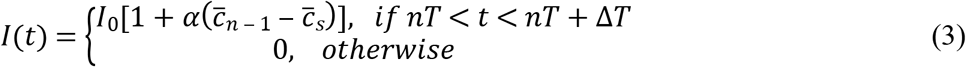

In Eq.3, *n* is the index of the beat number, *T* is the pacing period, Δ*T* is the pulse duration, and *α* is the feedback strength. 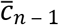 is the peak value of the spatial average of *c* (denoted as 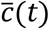 at the (n-1)^th^ beat. 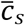 is the referenced value for the feedback. Here we define *α* > 0 as positive feedback (*α* < 0 as negative feedback), since in a single uncoupled FHN unit, a larger *c*_*n*‒1_ gives rise to a larger *I*(*t*), and thus a larger *c*_*n*_. We set *I*_0_=1.2, Δ*T*=0.5, and 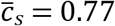.

### Excitation patterns without DGF

In the absence of DGF, i.e., *α* = 0 in Eq.3, a bifurcation from temporal P1 to P2 occurs as the pacing period T decreases (Fig.S2). We found that when T first passes through the bifurcation point, the system can only exhibit a spatially concordant P2 (Con-P2) pattern; as T decreases further, the system can exhibit a Con-P2 (Fig.2a) or a spatially discordant P2 (Dis-P2) pattern (Fig.2b), depending on initial conditions. It appears that the probability of forming a Dis-P2 pattern increases as the spatial heterogeneity of the initial condition increases (Fig.2c). Moreover, the Dis-P2 patterns are spatially random and selected by initial conditions. To quantify this property, we measured the spatial domain sizes (see Fig.2b for definition) from 2000 random trials for a given standard deviation of the spatial heterogeneity of the initial condition, and plotted the corresponding histogram (Fig.2d). It shows that the domain size can be any value as long as it is greater than a minimum domain size *l*_min_, i.e., the domain sizes distribute between *l*_min_ and *L*-*l*_min_. Because of this randomness in pattern selection, the corresponding histogram of the global P2 amplitude 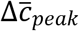 as defined in Fig.2a) also exhibits a continuous distribution (Fig.2e).

**Fig.2.**
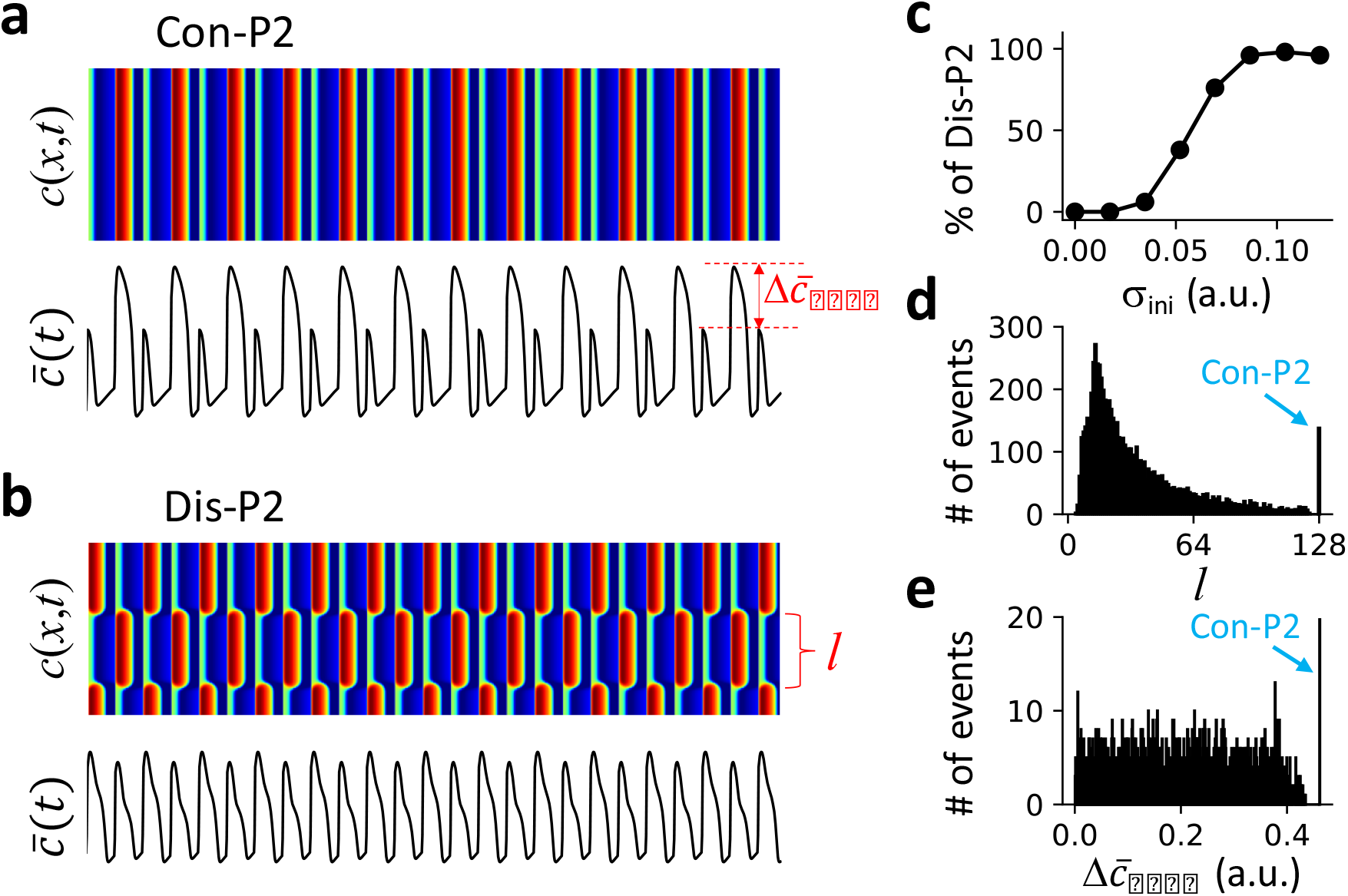
Excitation patterns and dynamics in a 1D array of coupled FHN units without DGF. The pacing period T=45 and system size L=128. **a**. An example of Con-P2 patterns and the corresponding global signal 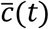. **b**. An example of Dis-P2 patterns with a different initial condition from panel a, and the corresponding 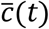. **c**. Percentage of Dis-P2 patterns versus the standard deviation (σ_*ini*_) of the random initial conditions. The random initial conditions were random spatial distributions of *w*(*j*), which was *w*(*j*) = *w*_0_ + *δξ*(*j*) (j ∈ {1,2,…,L}) with *ξ*(*j*) being a uniform random number drawn from 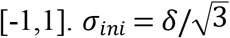. We set *w*_0_=0.5. We performed 100 trials for each σ_*ini*_ value in the plot. **d**. Histogram of domain size *l* (segment between two neighboring nodes, as indicated in panel b) from 2000 trials of random initial conditions with *δ* = 0.15. For each trial, 2000 beats were applied for the system to reach the steady state. The domain size was measured using the last two beats. **e**. Histogram of global P2 amplitude 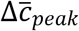, (difference between the peak values of two consecutive beats, as indicated in panel a) for the simulations in d. 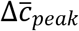 was measured using the last two beats.

### Effects of DGF on pattern selection and stability

To investigate the effects of DGF on the spatiotemporal pattern dynamics, we carried out simulations by scanning the pacing period T and DGF strength α (Fig.3a). There are four distinct regions: uniform P1 pattern (yellow), Con-P2 pattern only (cyan), Dis-P2 pattern only (black), and both concordant and discordant P2 (Con/Dis-P2) patterns (red). The blue curve is the stability boundary between P1 and P2 for a single uncoupled FHN unit. For *α* < 0, only uniform P1 and Con-P2 patterns were observed, independent of initial conditions. The uniform P1 and Con-P2 patterns are separated by the stability boundary (blue line) of the single uncoupled FHN unit, indicating that the dynamics in the 1D array is the same as in the single FHN unit. For *α* > 0, a transition from uniform P1 to Dis-P2 occurs as T decreases (from yellow to black), which is caused by a spatial-mode instability of the uniform P1 state. As T decreases further (red region), both Con-P2 and Dis-P2 patterns can occur depending on the initial conditions (Fig.3b).

**Fig.3.**
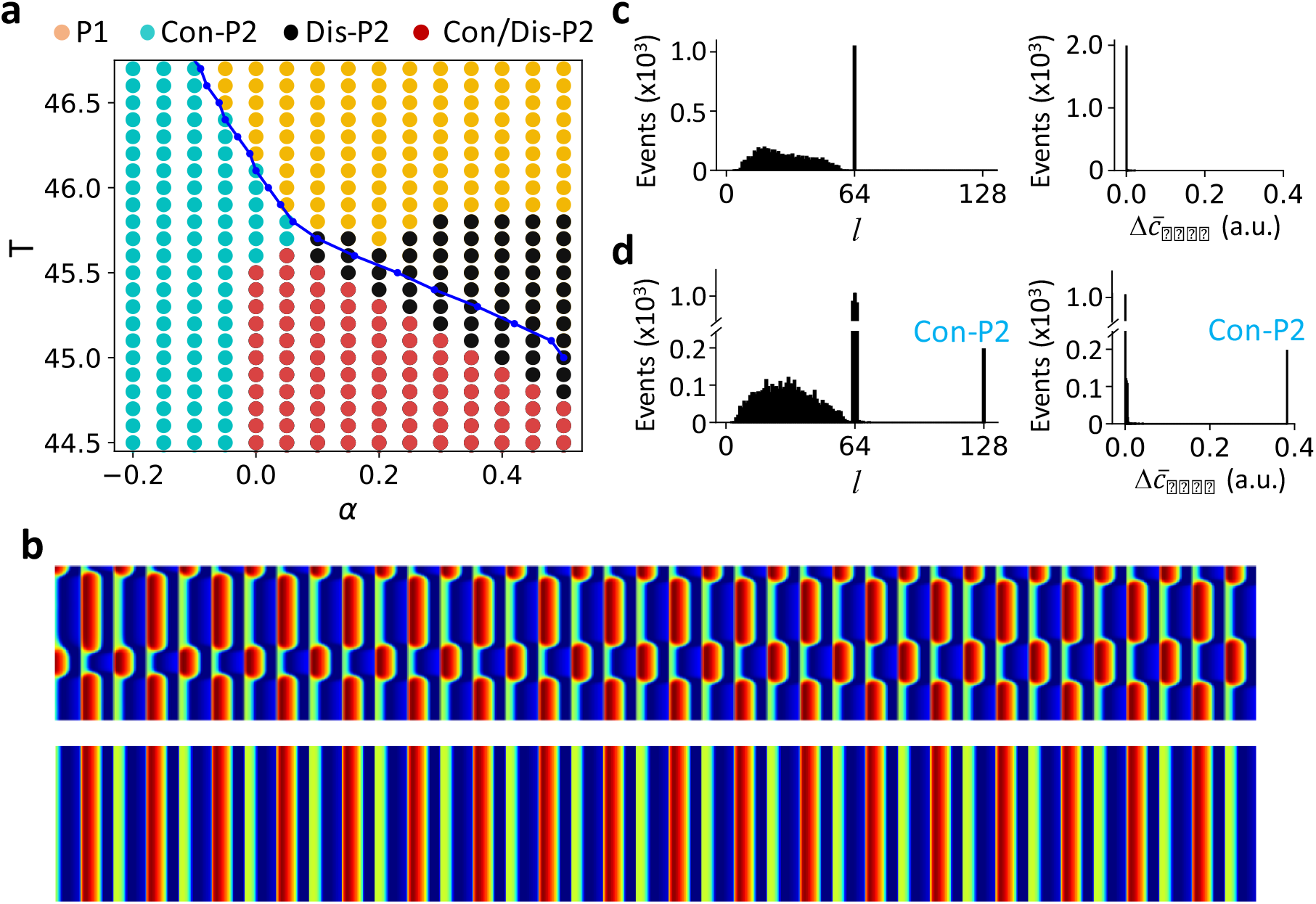
Excitation patterns and dynamics in a 1D array of coupled FHN units with DGF. **a**. Phase diagram of excitation dynamics. The blue line is the bifurcation boundary from P1 to P2 in a single uncoupled unit with DGF. Color dots mark the different behaviors in the 1D array: yellow—uniform P1; black—Dis-P2; cyan—Con-P2; and red—Con/Dis-P2. **b**. A Dis-P2 pattern (upper) and a Con=P2 pattern (lower) for *α* = 0.2 and T=45 obtained with two different initial conditions. **c**. Left, histogram of domain size *l* from 2000 trials. α=0.4 and T=45.5. The random initial conditions were set the same way as described in Fig.2 legend with δ=3. For each trial, 2000 beats were applied for the system to reach the steady state. Right, corresponding histogram of global P2 amplitude 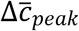 from the same simulations. **d**. Same as panel c but for α=0.2, T=45, and δ=0.09.

Furthermore, we performed the same statistical analysis as in the case of no DGF (Fig.2 d and e) for different regions. In the Dis-P2 only region (Fig.3c), the domain sizes distribute between 0 to *L*/2 (more accurately, the domain size can be *L*/2 and any value between *l*_min_ and *L*/2-*l*_min_), but 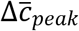 remains zero for all patterns. In the Con/Dis-P2 region (Fig.3d), the distributions are similar to those in Fig.3c except for the existence of the Con-P2 pattern. Similar to the case of no DGF (*α* = 0), the domain size distributions are continuous, indicating that the Dis-P2 patterns are spatially random (including the periodic ones) and depend on initial conditions. However, differing from the case of no DGF, the global signals of the Dis-P2 patterns are always P1 solutions i.e., 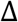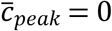 (Fig.3 c and d). Moreover, the maximum domain size of Dis-P2 patterns is *L*/2. This is because if there is a domain greater than *L*/2, the sum of all other domains must be smaller than *L*/2, and thus when the patterns reverse in the next beat, the global signal cannot be the same, violating the requirement of a global P1 solution.

Therefore, in the absence of DGF (*α* = 0), both Con-P2 and Dis-P2 patterns can occur, and the Dis-P2 patterns are spatially random. In the presence of DGF, only Con-P2 patterns can exist when the DGF is negative (*α* < 0). When the DGF is positive (*α* > 0), both Con-P2 and Dis-P2 patterns can exist depending on pacing period T and initial conditions. The Dis-P2 patterns are also spatially random but satisfy that the global signals are always P1 solutions.

### Ca^2+^ release patterns in a physiologically detailed ventricular myocyte model

To validate the spatiotemporal dynamics in a realistic biological system, we carried out simulations in a physiologically detailed 3D ventricular myocyte model (see Methods), which can well capture the spatiotemporal Ca^2+^ dynamics widely observed in experiments [29, 42–44]. The model undergoes a bifurcation from P1 to P2 (alternans) as the pacing period T decreases (Fig.S2). We investigated the subcellular Ca^2+^ release patterns under both AP clamp and free-running conditions. Under AP clamp (see Fig.S3 for the waveform used in this study), there is no DGF in the model. Under the free-running condition, DGF exists and its properties can be changed by altering ionic currents.

### Ca^2+^ release patterns under AP clamp

Under AP clamp, Ca^2+^ is decoupled with voltage. In the alternans regime (e.g., T=300 ms), both Con-P2 (Fig.4a) and Dis-P2 (Fig.4b) patterns occur in the cell depending on initial conditions. The probability of forming a Dis-P2 pattern increases as the spatial heterogeneity of initial conditions increases (Fig.4c). The Dis-P2 patterns are spatially random as indicated by the histograms of domain size (Fig.4d) and whole-cell alternans amplitude 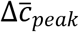 (Fig.4e). These behaviors are the same as those for the model of coupled FHN units without DGF (Fig.2), albeit some smearing in the histograms due to ion channel stochasticity.

**Fig.4.**
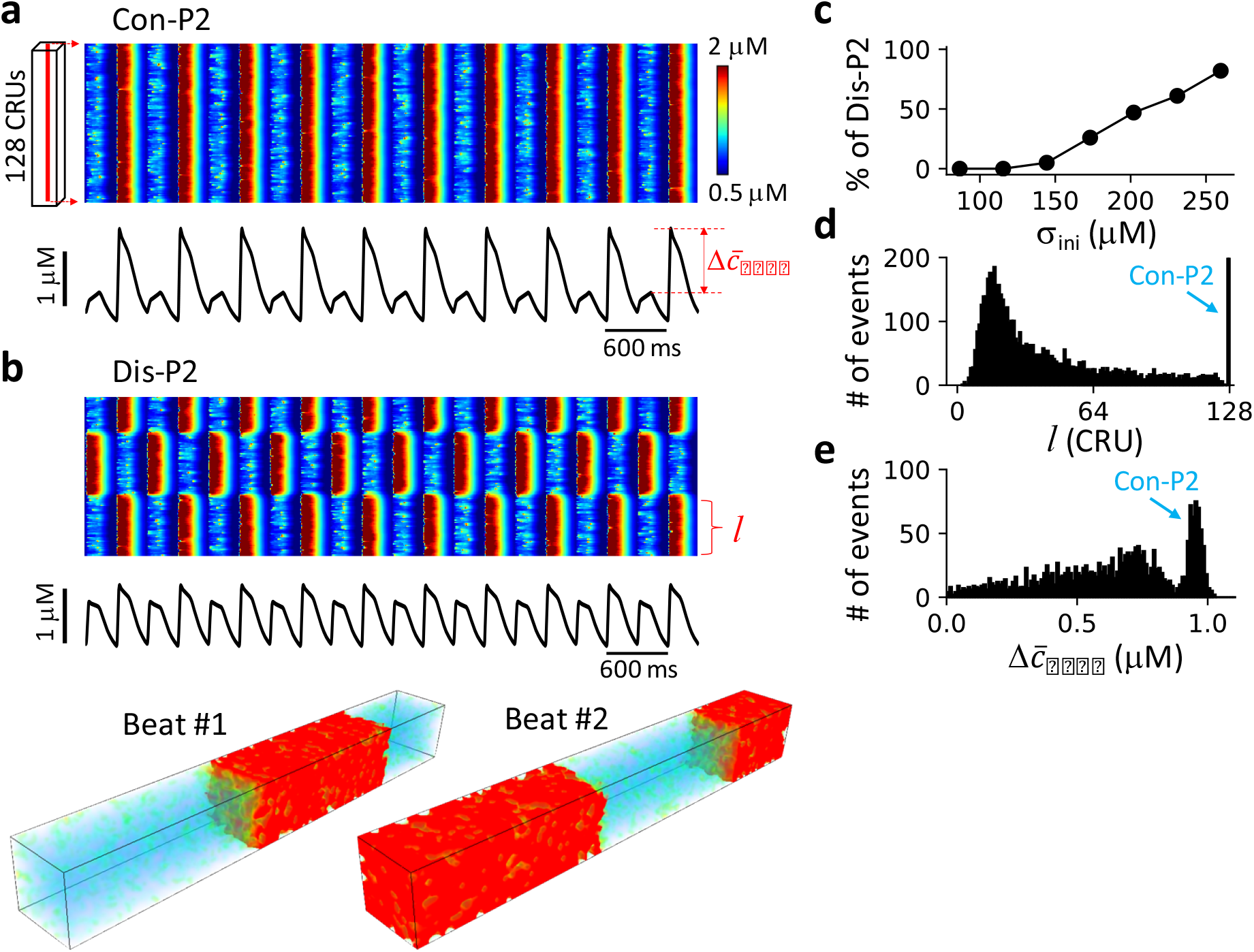
Ca^2+^ release patterns and dynamics in the 3D ventricular myocyte model under AP clamp. **a**. Upper panel shows a linescan (time-space plot) of cytosolic Ca^2+^ concentration showing a Con-P2 pattern. Lower panel shows the corresponding whole-cell Ca^2+^ transient. The recording line was in the center of the cell as indicated on the left. **b**. Same as panel a with a different random initial condition resulting in a Dis-P2 pattern. The middle panel is the corresponding whole-cell Ca^2+^ transient. The bottom panels are 3D views of Ca^2+^ from two consecutive beats. **c**. Percentage of Dis-P2 patterns versus the standard deviation (σ_*ini*_)of initial SR Ca^2+^ load. The random spatial distribution of the SR Ca^2*+*^ load was set as *Ca*_*SR*_(*j*) = *Ca*_0_ + Δ*Ca*_*SR*_ ⋅ *ξ*(*j*) (j ∈ {1,2,…,L}) with *ξ*(*j*) being a uniform random number in 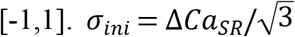. We set *Ca*_0_=500 μM. We performed 100 trials for each σ_*ini*_ value in the plot. **d**. Histogram of domain size *l* (as marked in panel b) with Δ*Ca*_*SR*_=450 μM. **e**. Histogram of global P2 amplitude Δ*c*_*peak*_ (as marked in panel a) from the same simulations in panel d. For panel d and e, 2000 trials were performed. For each trial, the cell was paced 2000 beats to reach the steady state. The domain size was computed using the last 50 beats to account for beat-to-beat variation (see SI for details) due to the intrinsic noise of ion channel stochasticity. 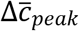 was measured using the last two beats. The pacing period T=300 ms.

### Ca^2+^ release pattern dynamics with positive and negative Ca^2+^-to-APD coupling

Under free running, however, Ca^2+^ is coupled with voltage, and changing Ca^2+^ may change APD. If increasing the Ca^2+^ transient amplitude results in a longer APD in the same beat, then it is called positive Ca^2+^-to-APD coupling, and the opposite is called negative Ca^2+^-to-APD coupling [45, 46]. To alter the coupling properties, we varied the maximum conductance of two Ca^2+^-dependent ionic currents in the model: the non-specific Ca^2+^-activated cation current (I_nsCa_) and the small conductance Ca^2+^-activated potassium current (I_SK_). Both currents increase as the Ca^2+^ transient amplitude increases. I_nsCa_ is an inward current such that an increase in Ca^2+^ transient prolongs APD, thereby enhancing positive Ca^2+^-to-APD coupling. I_SK_ is an outward current, which does the opposite, promoting negative Ca^2+^-to-APD coupling. We first investigated the effects of Ca^2+^-to-APD coupling properties on Ca^2+^ release patterns and then linked them to DGF.

We systematically explored the spatiotemporal dynamics by altering the pacing period T and the maximum conductance of the two currents, as summarized in Fig.5a. When the Ca^2+^-to-APD coupling is negative (large I_SK_), a transition from uniform P1 to Con-P2 patterns occurs as T decreases, and this transition occurs at a larger T value as the maximum I_SK_ conductance increases. When the coupling is positive (large I_nsCa_), a transition from uniform P1 to Dis-P2 patterns (yellow to black) occurs as T decreases. Under both coupling conditions, as T decreases further, the system enters the Con/Dis-P2 regime (red), in which both Con-P2 and Dis-P2 patterns can occur depending on initial conditions (Fig.5b). However, as T decreases even further, the Con/Dis-P2 regime switches into a Dis-P2 only regime when the Ca^2+^-to-APD coupling is negative (large I_SK_) and into a Con-P2 only region when the Ca^2+^-to-APD coupling is positive (large I_nsCa_). Therefore, for the same Ca^2+^-to-APD coupling, as T decreases, the spatiotemporal patterns change from Con-P2 only to Dis-P2 only through a Con/Dis-P2 region or in reverse order depending on the coupling properties.

**Fig.5.**
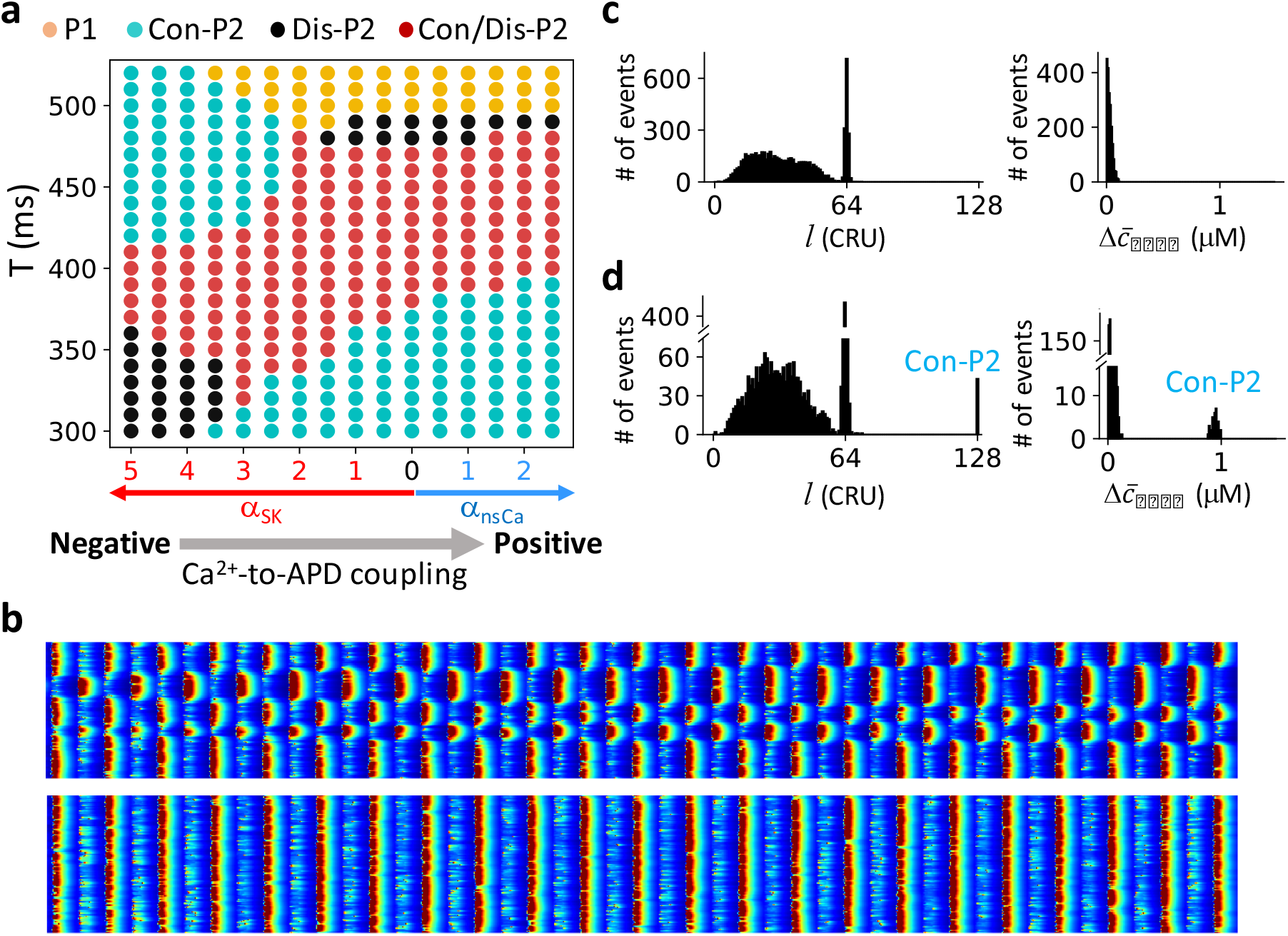
Ca^2+^ release pattern dynamics in the ventricular myocyte model with positive and negative Ca^2+^-to-APD coupling. **a**. Phase diagram of Ca^2+^ release dynamics versus pacing period and Ca^2+^-to-APD coupling properties. In this diagram, the x-axis is the fold increase of either *I*_nsCa_ (blue arrow) or *I*_SK_ (red arrow), and the y-axis is the pacing period T. Gray arrow indicates the change from negative to positive Ca^2+^-to-APD coupling. Same color codes of the pattern dynamics as in Fig.3a were used. **b**. A Dis-P2 pattern (upper) and a Con=P2 pattern (lower) for *α*_*SK*_ = 3.5 and T=350 ms obtained with two different initial conditions. **c**. Left: Histogram of domain size *l*. The pacing period T=330 ms, *α*_*SK*_ = 4.5, The random initial conditions were set the same way as in Fig.4 with Δ*Ca*_*SR*_=500 μM. 2000 trials were performed. For each trial, the cell was paced 2000 beats to reach the steady state. The domain size was computed using the last 50 beats. Right: 652 togram of global P2 amplitude Δ*c*_*peak*_ from the same simulations. 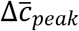 was measured using 653 the two beats. **d**. Same as panel c but T=350 ms and *α*_*SK*_ = 3.5.

To reveal the statistical properties of the Dis-P2 patterns, we show the histograms of domain sizes and the whole-cell alternans amplitude for a parameter point in the Dis-P2 region (Fig.5c) and a point in the Con/Dis-P2 region (Fig.5d). The domain size distributions for Dis-P2 patterns are continuous and the whole-cell alternans amplitudes always remain zero, indicating that the patterns are spatially random but always satisfying that the global signals are P1 solutions. These behaviors are the same as in the model of a coupled array of FHN units (Fig.3).

To link the spatiotemporal Ca^2+^ dynamics to the DGF properties, we performed an analysis to reveal the DGF properties and their relationship with the Ca^2+^-to-APD coupling properties. The details are described in the SI text and Fig.S4. This analysis shows that at fast pacing, positive (negative) Ca^2+^-to-APD coupling corresponds to negative (positive) DGF, which is mainly mediated via its effect on I_Ca_ recovery. At slow pacing rates, however, the relationships are reversed, and the DGF is primarily mediated via SR Ca^2+^ load since I_Ca_ fully recovers. Using the DGF properties, one can link the dynamics of the detailed physiological model to those of the generic model of coupled FHN units. In other words, the detailed model results validate the conclusion from the FHN model that only Con-P2 patterns can exist when the DGF is negative and both Con-P2 and Dis-P2 patterns can exist when the DGF is positive.

### Theoretical insights from a CML model

To reveal analytically the instabilities and bifurcations leading to the spatiotemporal dynamics, we used a CML model to describe the system. CML, as a generic model for investigating spatiotemporal dynamics of nonlinear systems, has been widely used [47, 48]. In a previous study [49], we developed a CML model to investigate the spatiotemporal APD dynamics in cardiac tissue. Here we modified the 1D array CML model by adding a DGF term. The governing equation is,

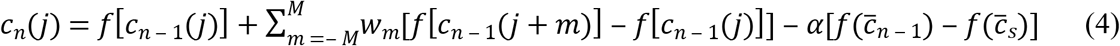

where *n* is the temporal index and *j* the spatial index. *c*_*n*_(*j*) describes the peak signal in the *j*^th^ lattice of the *n*^th^ beat. *f* is the map function: 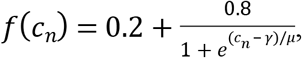 in which γ and *μ* determine the midpoint and the slope of the curve, respectively. M is the coupling length, and w_m_ is the coupling strength described by a Gaussian function: 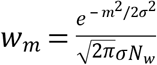 in which *N*_*w*_ is the normalization constant. 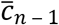 is the spatial average of *c*_*n*‒1_(*j*), and 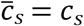 satisfying *c*_*s*_ = *f*(*c*_*s*_). We set *μ*=0.1, M=15, and σ=3.

Note that in Eq.4, instead of using a linear feedback term: 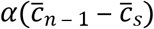, we used a nonlinear term with the map function *f*, i.e., 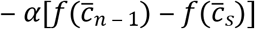, to maintain the convergence of iteration. The negative sign was used because *f* is a decreasing function (*f*’ < 0). Linearization of this nonlinear term around the uniform steady-state gives rise to a term proportional to 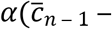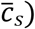, and thus *α* > 0 corresponds to positive feedback, the same as in the FHN model.

1. *Stability of a single uncoupled unit*. For a single uncoupled unit, the map equation with DGF becomes

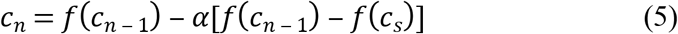 The stability of the steady-state solution is determined by the eigenvalue,

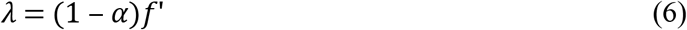

where 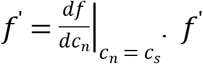 is independent of *α* since the steady state is independent of *α*. Eq.6 indicates that *α* < 0 destabilizes the steady state, and *α* > 0 stabilizes the steady state. The stability boundary is shown as the dashed line in Fig.6a.
2. *Stability of the spatially uniform P1 state*. The spatially uniform P1 state (see Stability analyses of the CML model in *SI*) is determined by the eigenvalues:

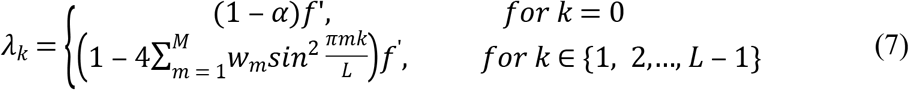

in which *k* is the wave number of the Fourier mode (*λ*_*k*_ vs. *k* for different *α* values are shown in Fig.S5). The spatially uniform P1 state is stable when |*λ*_*k*_| < 1 for any *k*. The stability of the 0-mode is the same as that of a single uncoupled unit. Since *λ*_*k*_ for *k* > 0 in Eq.7 does not depend on *α*, then the feedback has no effects on the stability of the uniform P1 state for non-zero mode. Because of this, the stability boundary separating uniform P1 from Dis-P2 appears to be a horizontal line independent of α (Fig.6a, solid).
3. *Stability of the Con-P2 state*. Following the same procedure as for the uniform P1 state, we obtained the eigenvalues for the spatially uniform P2 state as

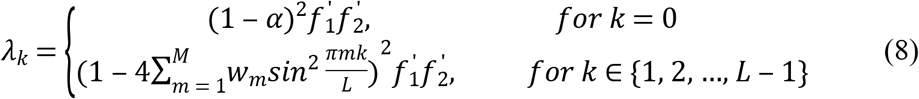

where 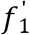 and 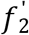 are the two derivatives of *f* at the P2 solution of Eq.5. Since the P2 solution depends on α, 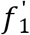 and 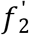 are functions of α. Therefore, the stability boundary also depends on α (Fig.6a, dashed-dotted).
4. *Stability of the Dis-P2 states*. The stability of the Dis-P2 states cannot be analytically obtained. We used numerical simulations of the CML model (Eq.4) to determine the stability boundary (Fig.6a, dotted). No stable Dis-P2 patterns were obtained on the left side of the dotted line.

**Fig.6.**
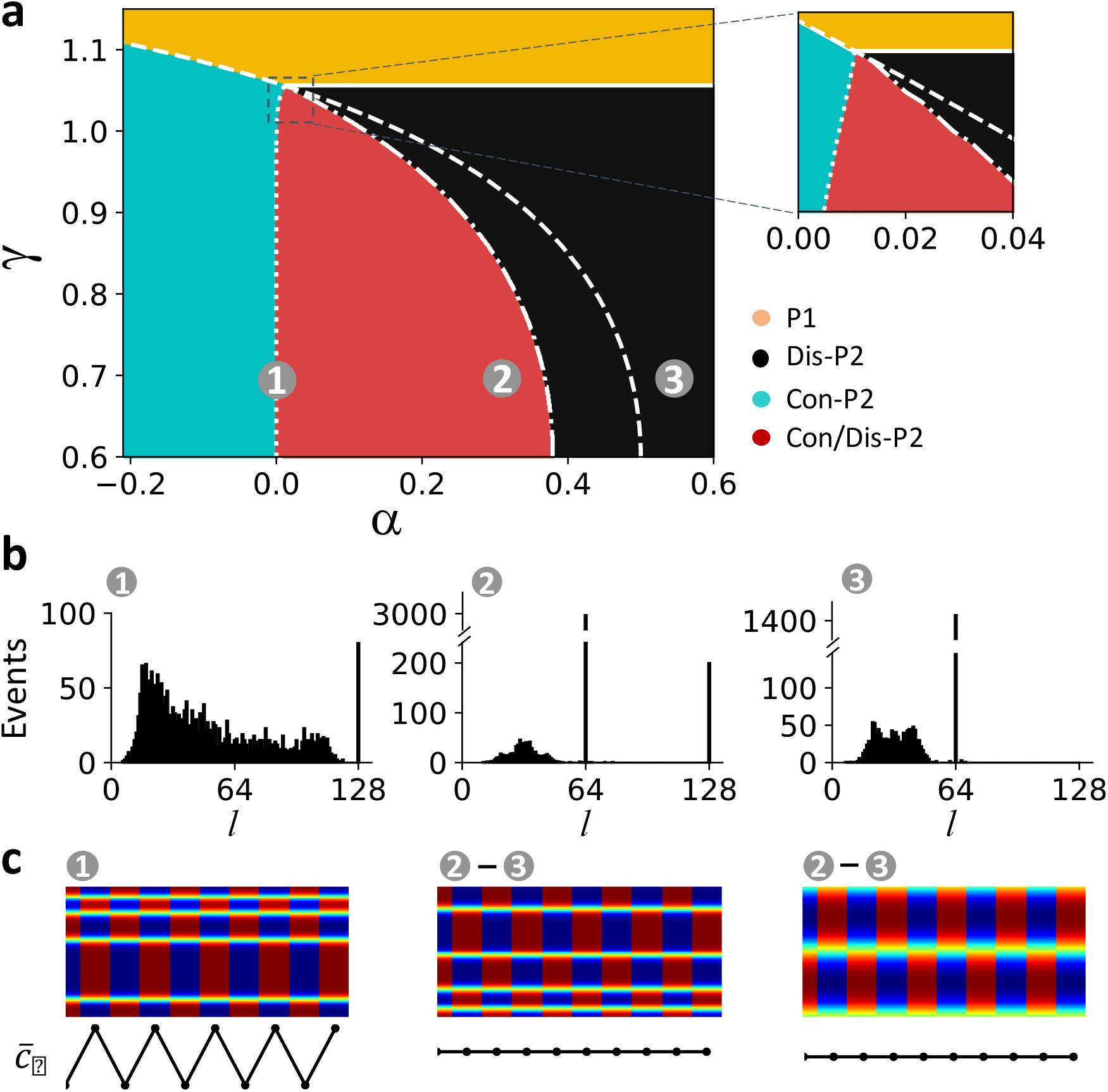
Bifurcations and spatiotemporal dynamics in the CML model. **a**. Phase diagram in the α-γ space showing stability boundaries and spatiotemporal dynamics of the CML model. The same color codes as in Fig.3a and Fig.5a were used. The solid line is the stability boundary of uniform P1 determined by Eq.7. The dashed line is the stability boundary of P1 in a single uncoupled unit determined by Eq.6. The dash-dot line is the stability boundary of Con-P2 determined by Eq.8. The vertical dotted line is the stability boundary of Dis-P2 determined by numerical simulations of the CML model. Inset is the blowup of the marked region showing that all the stability boundaries meet at a common point. **b**. Histograms of domain size at different locations in the phase diagram marked by numbers. The coordinates (α, γ) from location 1 to 3 are: (0, 0.7), (0.3, 0.7), and (0.55, 0.7). **c**. Sample spatiotemporal patterns (top) and the corresponding global signals (bottom) from different regions of the phase diagram. Number ranges above each pattern indicate the locations in the phase diagram where the specific pattern can be seen.

Spatiotemporal dynamics via numerical simulations of the CML model are also shown in Fig.6a, which are colored the same way as in Fig.3a and Fig.5a. The Dis-P2 only region exists between the uniform P1 stability boundary (solid line) and the Con-P2 stability boundary (dash-dotted line). The Con/Dis-P2 region exists between the Con-P2 stability boundary (dashed-dotted line) and the Dis-P2 stability boundary (dotted line). The Con-P2 only region exists between the uniform P1 stability boundary (dashed line) and the Dis-P2 stability boundary (dotted line). Note that the dotted line is almost identical to *α* = 0 except at the vicinity where all phases meet (inset in Fig.6a), indicating that stable Dis-P2 patterns can only exist when *α* > 0. Histograms of domain size and example spatial patterns from three locations marked in Fig.6a are plotted (Fig.6 b and c). The structure of the phase diagram and the statistical properties of spatial patterns of the CML model match well with those of the generic FHN model and the ventricular myocyte model.

## Discussion

We investigated the roles of DGF in the genesis, selection, and stability of spatiotemporal patterns in periodically-paced excitable media. We used a multi-scale approach in which three models with different complexities were utilized. The dynamical behaviors are well conserved in the three scales of models, and the CML model reveals the dynamical mechanisms. Our major findings are as follows:

1. In the absence of DGF, both Con-P2 and Dis-P2 can occur depending on the pacing period and initial conditions. The Dis-P2 patterns are spatially random, determined by the initial conditions. The global signal (the spatial average) is a temporal P2 solution (alternans) with the alternans amplitude being randomly distributed between zero and the maximum amplitude (Con-P2).
2. In the presence of DGF, the pattern dynamics are determined by the sign of the DGF. When the DGF is negative, only Con-P2 patterns can exist, no spatial mode instabilities emerge, and all the Dis-P2 solutions are unstable. When the DGF is positive, both Con-P2 and Dis-P2 patterns can occur, depending on the pacing period and initial conditions. The Dis-P2 patterns are also spatially random but must satisfy that the global signals are temporal P1 solutions (no temporal alternans).
3. Bifurcation analyses of the CML model reveal the spatial-mode instabilities leading to the spatiotemporal patterns.
4. By linking the Ca^2+^-to-APD coupling properties to the DGF properties, we have shown that the spatiotemporal pattern dynamics of Ca^2+^ release in cardiac myocytes agree very well with the findings in the simple models, validating the theoretical predictions in a realistic system.

Therefore, our simulations and theoretical analyses reveal the underlying dynamical mechanisms and roles of DGF in the genesis, selection, and stability of spatiotemporal patterns in periodically-paced excitable media. The uniqueness of the conclusions drawn from the multi-scale modeling approach implies that the insights obtained in this study may apply to many excitable as well as oscillatory biological media. Here we discuss two examples below.

### A unified theory for subcellular Ca^2+^ alternans dynamics in cardiac myocytes

As shown in this study, the subcellular Ca^2+^ alternans dynamic of the ventricular myocyte model agree well with those of the simplified models, indicating that the generic mechanisms of pattern formation and selection are also applicable to Ca^2+^ alternans dynamics in cardiac myocytes. Both spatially concordant and discordant Ca^2+^ alternans (Con-P2 and Dis-P2 patterns) have been observed experimentally in cardiac myocyte [31–33, 50]. Shiferaw and Karma [51] developed a theory showing that a Turing instability caused by negative Ca^2+^-to-APD coupling is responsible for the formation of Dis-P2 patterns. A direct experimental demonstration of this theory was carried out by Gaeta et al. [33], who developed a method that could change the sign of Ca^2+^-to-APD coupling. However, Dis-P2 patterns have also been observed experimentally under voltage-clamp [32, 50] and free-running conditions without showing negative coupling [52]. Furthermore, previous simulation studies [43, 53] and this study have also shown that Con-P2 patterns can occur under negative Ca^2+^-to-APD coupling, and Dis-P2 patterns can occur under positive Ca^2+^-to-APD coupling and voltage-clamp conditions. These complex Ca^2+^ release behaviors cannot be well explained by the Turing instability mechanism alone. On the other hand, our study unifies the complex subcellular Ca^2+^ alternans dynamics under a single theoretical framework of DGF, providing a general mechanistic understanding of the subcellular Ca^2+^ alternans dynamics.

### Links to pattern dynamics in oscillatory media with DGF

Our study focused on the roles of DGF in pattern formation and stability in periodically-paced excitable media. In a previous study in oscillatory chemical reaction experiments, Kim et al. [17] showed DGF caused clustering patterns similar to the Dis-P2 patterns in this study. Their observations were also demonstrated in computer simulations [19]. Since, in their studies, the DGF is an externally controlled signal, the delay time is a variable parameter. However, the DGF is intrinsic in the excitable biological media we investigated, and the delay time is simply the excitation period. Because of this, we can represent the system with a CML model that is able to capture the dynamics and the underlying bifurcations accurately. Since an excitable medium can become an oscillatory medium, the theories from our study may provide mechanistic insights into pattern dynamics of oscillatory media, such as clustered firings of oscillatory neural systems.

## Methods

The present study involved three mathematical models at different levels of complexity. The model of a coupled array of FHN units and the 1D CML model are described in the Result section. A brief summary of the 3D ventricular cell model and numerical methods is given below.

### The 3D ventricular cell model

The ventricular cell model has been described in detail in our previous studies [43, 44], similar to other previous models [54–58]. Here we give a brief description of the model. The 3D cell model consists of 128 × 16 × 16 CRUs. Each CRU includes five sub-compartments: bulk cytosol, submembrane, dyad, junctional SR and network SR. The volumes of these sub-compartments are based on experimental data. The Ca^2+^ within a CRU cycles through these sub-compartments via diffusion, buffering/unbuffering, SR release and SERCA pump. The flow of Ca^2+^ between CRUs is via diffusion in the cytosol, submembrane and network SR. The exchange of Ca^2+^ between intracellular and extracellular space is regulated by I_Ca_ and Na^+^-Ca^2+^ exchanger (NCX).

We added two new currents, *I*_*nsCa*_ and *I*_*SK*_, to the model for altering Ca^2+^-to-APD coupling. The *I*_*nsCa*_ formulation was adopted from the 1994 Luo and Rudy model [59] with the following parameter changes: P_ns(Ca)_=1.75 × 10^−7^ and K_m,ns(Ca)_=1.5 μM. *I*_*SK*_ was formulated based on Komendantov et al [60] as follows:

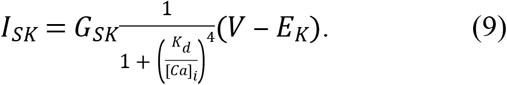

The differential equation for voltage is then

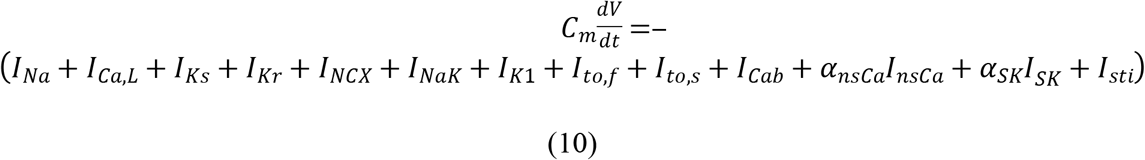

where *C*_*m*_=1 *μF*/*cm*^2^ is the membrane capacitance. α_nsCa_ and α_SK_ are the parameters controlling the maximum conductance of *I*_*nsCa*_ and *I*_*SK*_, respectively. *I*_*sti*_ is the stimulus current density which is a square pulse with the amplitude −80 μA/μF and the duration 0.5 ms.

### Computer simulations and algorithms

The model of a coupled array of FHN units and the CML model were programmed with Python 3, and the corresponding simulations were carried out on our cluster with 24 Intel® Xeon® CPUs. The 3D ventricular cell model was programmed with CUDA C++, and the corresponding simulations were carried out on Nvidia Tesla K20c, K80, and GTX 1080 Ti GPU cards. The detailed algorithms for detecting spatiotemporal excitation patterns in this study are described in the ****SI**** text and Fig.S6.

## Acknowledgments

This work was supported by grants from National Institutes of Health R01 HL133294 and R01 HL134709.

## Supporting information

**SI Text. Linking Ca^2+^-to-APD coupling properties to DGF properties.**

**SI Text. Stability analysis of the CML model.**

**SI Text. Automatic detection algorithms for spatiotemporal excitation patterns.**

**SI Text. Boundary between discordant P2 and uniform P2 in the CML model.**

**S1 Fig. Examples of spatiotemporal Ca^2+^ release dynamics in excitable biological systems.**

**S2 Fig. Bifurcation diagrams for the FHN, detailed ventricular myocyte and CML models.**

**S3 Fig. Time trace of membrane voltage used in the AP clamp protocol for the detailed cell model.**

**S4 Fig. Relationship between Ca^2+^-to-APD coupling and DGF.**

**S5 Fig. *λ*_*k*_ vs. the wave number***k* **for the 1D CML model under the P1 regime.**

**S6 Fig. Illustration of domain size detection by the pattern recognition algorithm.**

**S1 Table. Parameters of nsCa and SK currents.**

